# A semi-automated method for measuring xylem vessel length distribution

**DOI:** 10.1101/2020.08.04.234575

**Authors:** Luciano Pereira, Marcela T. Miranda, Gabriel S. Pires, Vinícius S. Pacheco, Xinyi Guan, Lucian Kaack, Eduardo C. Machado, Steven Jansen, Melvin T. Tyree, Rafael V. Ribeiro

## Abstract

Knowledge about the length of xylem vessels is essential to understand water transport in plants because these multicellular units show a 100-fold variation, from less than a centimeter to many meters. However, the available methods to estimate vessel length distribution (VLD) are excessively time consuming and do not allow large and in-depth surveys. Here, we describe a semi-automated method to measure VLD using an automated Pneumatron device. Gas conductivity of a xylem tissue with a certain length is estimated in a straightforward and precise way with the Pneumatron in a way theoretically similar to the air-injection method. The method presented enables fast and easy measurements using multiple devices simultaneously (>50 samples day-1), which is a significant advantage. Here, the apparatus is described in detail as well as how measurements are taken. We also present the software and an R-script for data analysis. The method described represents an important contribution to studies on plant hydraulic architecture and can improve our understanding about the role of VLD in plant performance under varying water availability.

## 1 Introduction

Vessels in angiosperm xylem are formed by stacking of individual vessel elements, which are typically jointed near their tips by a single or multiple perforation (Medeiros et al. 2019). Whereas individual vessel elements are less than one millimeter, the total vessel length may vary from one centimeter to more than 10 meters (Greenidge 1952). Besides the connections via perforation plates, which are relatively large openings (>2 μm) through the secondary and primary cell walls, vessels are interconnected by openings in the secondary wall that are sealed by nano-scale perforations in the primary walls of pits (Kaack et al. 2019). The large variation in vessel length is also related to considerable variation in vessel width, which varies from ca. 10 µm to more than 300 µm (Cai and Tyree 2010; Pan et al. 2015; Liu et al. 2018). Moreover, with many vessels being very short, and a smaller proportion relatively long, measuring the vessel length distribution in xylem has been challenging for a long time (Williamson and Milburn 2017; Link et al. 2018).

Among the available methods, the silicone-injection method to study vessel length distribution (VLD) in a stem/root segment is considerably time-consuming (Cai and Tyree 2014). This method consists of perfusing silicone with a particular dye into plant samples. The silicon typically does not pass through the tiny nano-scale pores of the pit membranes between adjacent vessels (Zhang et al. 2020). Then, the number of vessels with lumina filled with silicone is measured with the aid of a microscope by examining several transverse sections at several distances from the injection point. When using the silicone-injection method, one to three weeks are needed to measure five branches (Pan et al. 2015). The time-consuming nature of this protocol may be the reason why relatively few papers on plant hydraulics include vessel length distribution data (Jacobsen et al. 2012). Alternatively, measurements on the VLD were greatly facilitated by the gas-conductivity method (Cohen et al. 2003), and the more simplified protocol proposed by Wang et al. (2014). This method evaluates VLD by injecting gas at one side of a plant sample and measuring the gas conductivity through a segment with a particular length, with the gas outlet volume being collected over a certain period of time. Then, the segment is shortened, and the gas conductivity is successively measured until all vessels are open (Wang et al. 2014). Although this method is faster (e.g. few hours), the simplified gas-conductivity method is still less frequently used as compared to the silicone-injection method, and as far as we know, there is no report in which this method was used for large and in-depth surveys, possibly due to the need of laboratory equipment and accessories (e.g. balance, compressor and flasks), and the handling of gas.

Inspired by the development of a recent device to measure embolism formation in dehydrating xylem (Pereira et al. 2020), we present a semi-automated method to measure VLD using the Pneumatron. This instrument is based on the Pneumatic method (Pereira et al. 2016; Bittencourt et al. 2018; Jansen et al. 2020), which measures the amount of gas that can be extracted from stems (Pereira et al. 2016; Zhang et al. 2018), roots (Wu et al. 2020), or leaves (Pereira et al. 2020). By measuring pressure changes inside a fixed tube volume over time, gas conductivity is calculated by extracting gas from plant xylem under sub-atmospheric pressure. This approach was tested here to measure VLD, but instead of small volumes of gas from embolized vessels, we measured the gas conductivity through open vessels in branch segments with variable length. Under these conditions, the gas flow varies linearly with pressure change and can be directly related to hydraulic conductivity (Cohen et al. 2003). Although the method principle is similar to one reported by Cohen at al. (2003), we tested if sub-atmospheric and variable pressure created by the Pneumatron can be used in a similar way as the positive and constant pressure originally proposed (Cohen et al. 2003).

Besides the description of the simple apparatus and some VLD estimations for *Citrus limonia, Eucalyptus camaldulensis, Eugenia uniflora, Olea europaea* and *Persea americana*, we also present the specific software that needs to be uploaded to the Pneumatron and a R-script for speeding up the data analysis and VLD estimations.

## 2 Materials and Methods

### 2.1 Plant material and apparatus

We used stems taken from mature field-grown trees of *Citrus limonia* Osbeck, *Eucalyptus camaldulensis* Dehnh., *Eugenia uniflora* L., *Olea europaea* L., and *Persea americana* Mill. grown at Campinas SP, Brazil (22°54’23"S, 47°3’42"W, 600 m a.s.l). Although the measurements were straightforward, we only studied a few species to demonstrate this new method. Hence, a systematic study on vessel length distribution based on a large dataset is beyond the scope of this methodological paper. The stems had a diameter of *c.a*. 10 mm and no lateral ramification. The cut surfaces were shaved with a sharp razor blade to avoid obstruction of vessels, and the proximal end was connected to the apparatus using a flexible silicone tube, clamp, luer adaptor, and rigid tubbing, following the same procedure described for estimating embolism vulnerability curves (Bittencourt et al. 2018; Pereira et al. 2020). A rigid tube was connected to a 100 mL flask reservoir, which was then connected to the Pneumatron (Fig. 1). A smaller reservoir volume of 3 mL was used for *Citrus limonia* and *Eucalyptus camaldulensis* measurements for testing the method precision (see section 2.3).

**Fig. 1.**
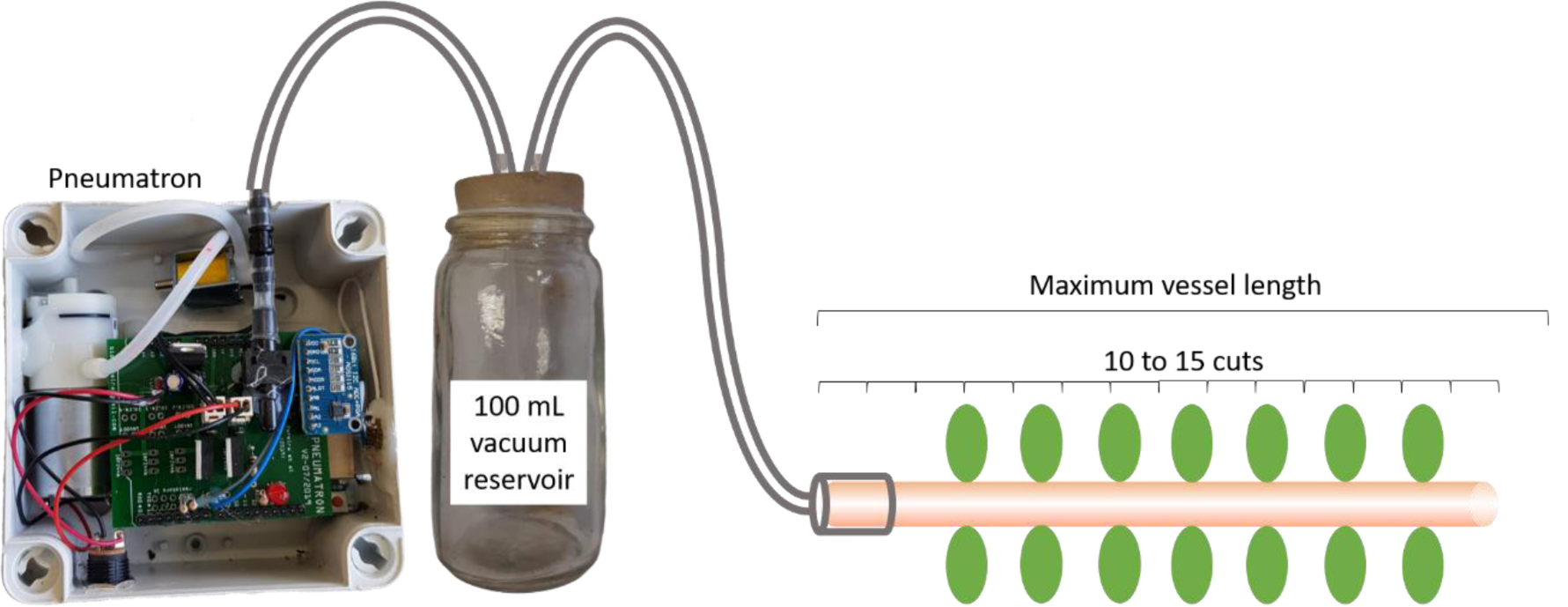
Apparatus setup for measuring the vessel length distribution with the Pneumatron. Stem segments shorter than the maximum vessel length were used and successive cuts (10 to 15) were made while gas conductivities were measured.

A complete description of the Pneumatron can be found in Pereira et al. (2020). For estimating VLD, the hardware was not changed, and it was similar to the device originally described for measuring embolism. Briefly, the Pneumatron included the following operational steps: a vacuum pump and solenoid valve were turned on to decrease the pressure inside the discharge tube down to 40 kPa (absolute pressure); at this pressure, both the pump and solenoid valve were turned off, and pressure changes monitored for 150 seconds. During this time, the pressure was kept between 40 and 50 kPa by turning on the pump automatically, of which the frequency depended on the gas flow (Fig. 2). The pressure variation within 150 seconds was recorded in a memory card and used for calculating the gas conductivity (see section 2.2). We did not consider the pressure values recorded immediately after turning on the vacuum pump as it resulted in a decrease of the absolute pressure.

**Fig. 2.**
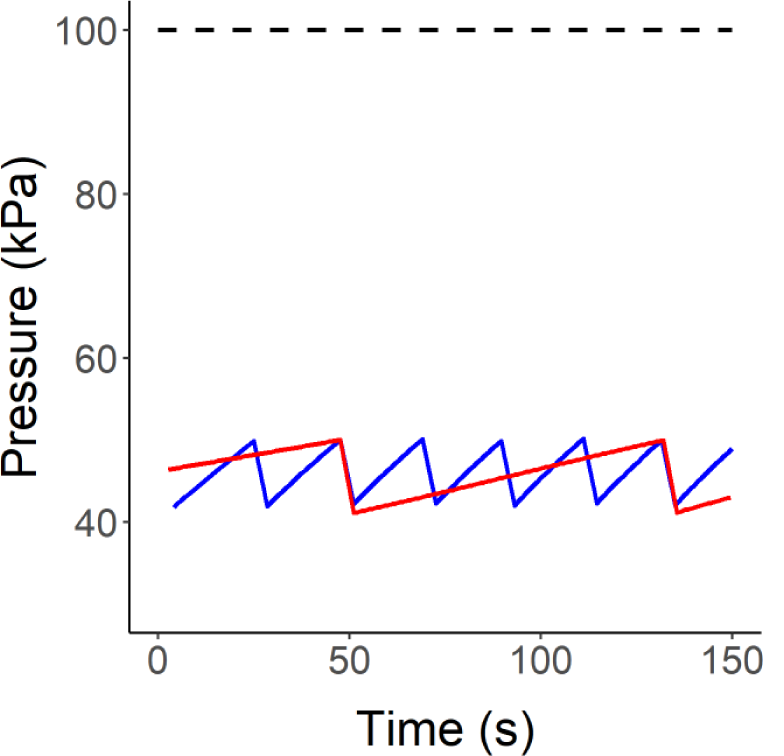
Absolute pressure change during the measurements of air flow using the Pneumatron. Measurements were taken at long (0.3 m, in red) and short (0.15 m, in blue) intervals of the same stem (diameter = 8.5 mm) of *Eugenia uniflora*. The horizontal dashed line represents the atmospheric pressure.

The absolute pressure difference between both segment ends (50 kPa) was similar to the above-atmospheric pressure used in the original method (60 kPa, Cohen et al. 2003). Air-seeding in intact vessels is supposed to occur with a much higher difference of pressure (> 1 MPa), considering the nanopores size in the pit membranes (Kaack et al. 2019). Thus, intact vessels are kept water filled and are not conductive to gas until they are cut open.

### 2.2 Measuring vessel length

To measure VLD, firstly the maximum vessel length (MVL) of a species was measured to estimate the initial stem length, and secondly the actual measurements were taken while successively cutting off the sample stem (Fig. 1). The initial stem length was defined for each species using segments shorter than the MVL, which means that at least some long vessels were cut open when the measurements were started. The MVL was previously estimated for each species by using the air-injection method (Greenidge 1952).

Based on Cohen et al. (2003), unbranched stem segments were long enough to obtain 10 to 15 measurements, i.e. 10 to 15 cuts, which considerably increased the number of open vessels. This is needed because measurements must be taken at the exponential increase of gas conductivity as a function of length for VLD estimations (Cohen et al. 2003). Data collected before such exponential increase should be disregarded.

The stem segment was successively shortened, and the respective gas conductivity measured. Depending on the species and the MVL, the segments were cut every 1, 3, or 5 cm, and the stem length measured. The cut surface was shaved with a razor blade after each cut and after *ca*. 10 to 15 cuts, the data saved on the Pneumatron memory card were transferred to a computer and analyzed using an R-script (Supplementary Material). For checking data stability during the measuring time, the pressure was recorded every 0.5 second and the gas flow was calculated for every 1 second for samples of *Eugenia uniflora, Olea europaea* and *Persea americana*. We noticed that gas conductivity was already stable after 50 seconds for these three species (Fig. S1). The initial instability is probably due to water extraction from the recently cut open vessels, although theoretically, this extraction would occur in the first few seconds (Cohen et al. 2003). Thus, to ensure measurement stability, the VLD estimation considered the gas conductivity averaged over the last 30 seconds (between 120 and 150 seconds) for all species.

Firstly, we calculated the amount of gas discharged per second (GD, mol s-1) inside the Pneumatron discharge tube according to the equation 1:

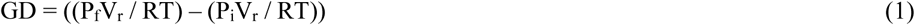

Where P_i_ and P_f_ are the initial and final pressures (within one second), V_r_ is the reservoir volume (in m3), R is the gas constant (8.3144621 J mol-1 K-1) and T is the air temperature (in K).

The gas conductivity (C, mol m s-1 Pa-1) was calculated according to Pan et al. (2015) as:

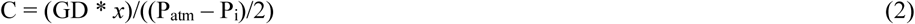

Where *x* is the stem length (in m) and P_atm_ is the atmospheric pressure (in Pa).

The slope (λ_v_) of the ln(C) *vs. x* was then estimated, and the number of open vessels (N) calculated as follows:

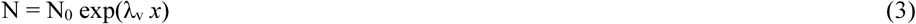

Where N_0_ is the number of open vessels when *x* = 0, that is, the intercept of the ln(C) *vs. x*.

Then, the probability distribution function (P_x_), the most common vessel length (L_mode_), and the mean vessel length (L_mean_) were calculated (Cai and Tyree 2014), as shown below:

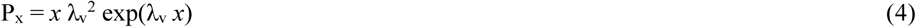

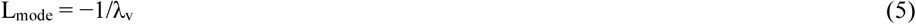

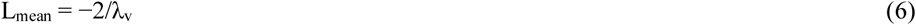

### 2.3 Measurement precision and reservoir volume

As shown in equation 1, a precise measurement of gas flow is directly dependent on the reservoir volume as well as the pressure reading. For each reservoir volume, it is possible to measure a determined range of gas flow. The minimum gas flow that can be detected is determined by the pressure sensor linearity (0.25% or 0.25 kPa considering the full scale of the sensor as 100 kPa), that is, the gas flow that can induce 0.25 kPa of pressure change. On the other hand, the maximum gas flow that can be measured depends on the pressure range that is controlled by the Pneumatron (40 to 50 kPa), and a minimum period to increase the pressure from 40 to 50 kPa. Here, we considered 3 seconds to obtain at least three conductivity measurements. From those conditions described above, we calculated the linear relationships (Fig. 3) for the maximum (slope = 1.36759E-06) and minimum (slope = 1.02569E-07) reservoir volumes that can be used for measuring gas flow. Thus, we estimated the minimum and maximum reservoir volumes that could be used for each flow measured as described in section 2.2. It was possible to check if the reservoir volume could be used for measuring the gas flow range as accurately as possible for each sample.

**Fig. 3.**
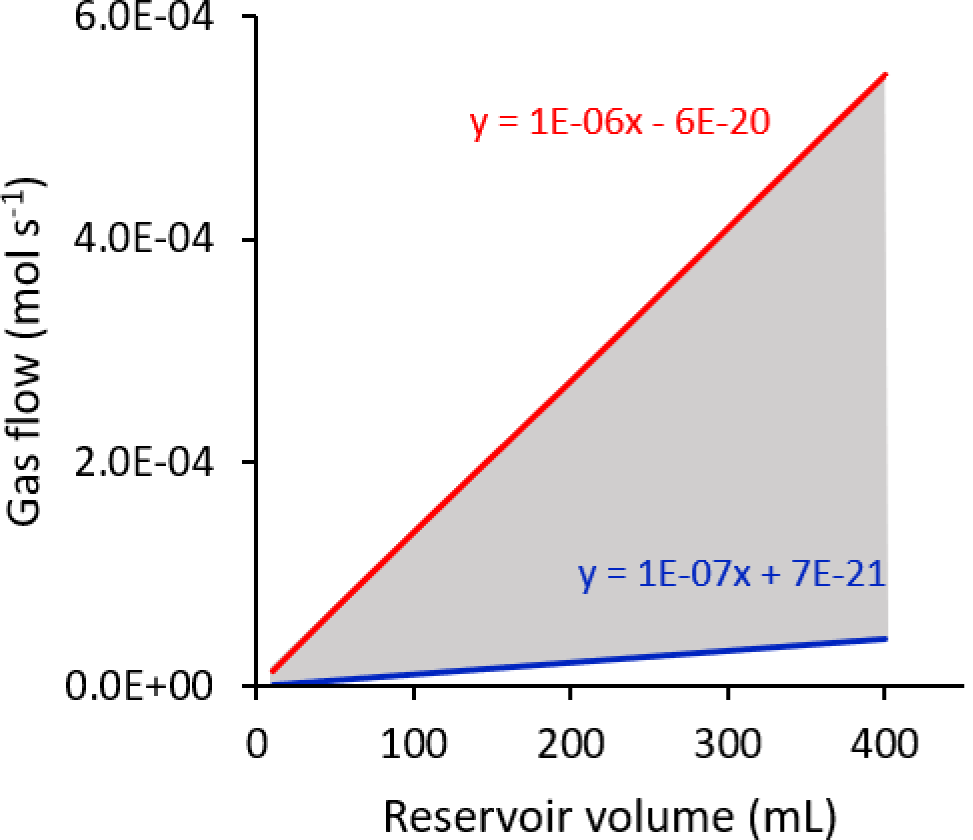
Maximum (red) and minimum (blue) gas flow that can be detected depending on the reservoir volume used for the measurements. To estimate the minimum gas flow, we considered the pressure sensor linearity (0.25%), and the fastest change (3 seconds) of the pressure range that is used for the measurements (between 40 and 50 kPa, see Materials and Methods for details) for determining the maximum gas flow. The grey area represents the range of gas flow that can be measured for a given reservoir volume.

## 3 Results

Gas discharged was exponentially related to stem length for all species (Figs. 4a and S2a) and there was a high correlation between stem segment length and the natural log of gas conductivity (*r2* > 0.97, Figs. 4b and S2b). When the *r2*-value was lower than 0.97, the sample was disregarded for the VLD estimation as it may indicate a measurement error due to leakage (Cohen et al. 2003).

**Fig. 4.**
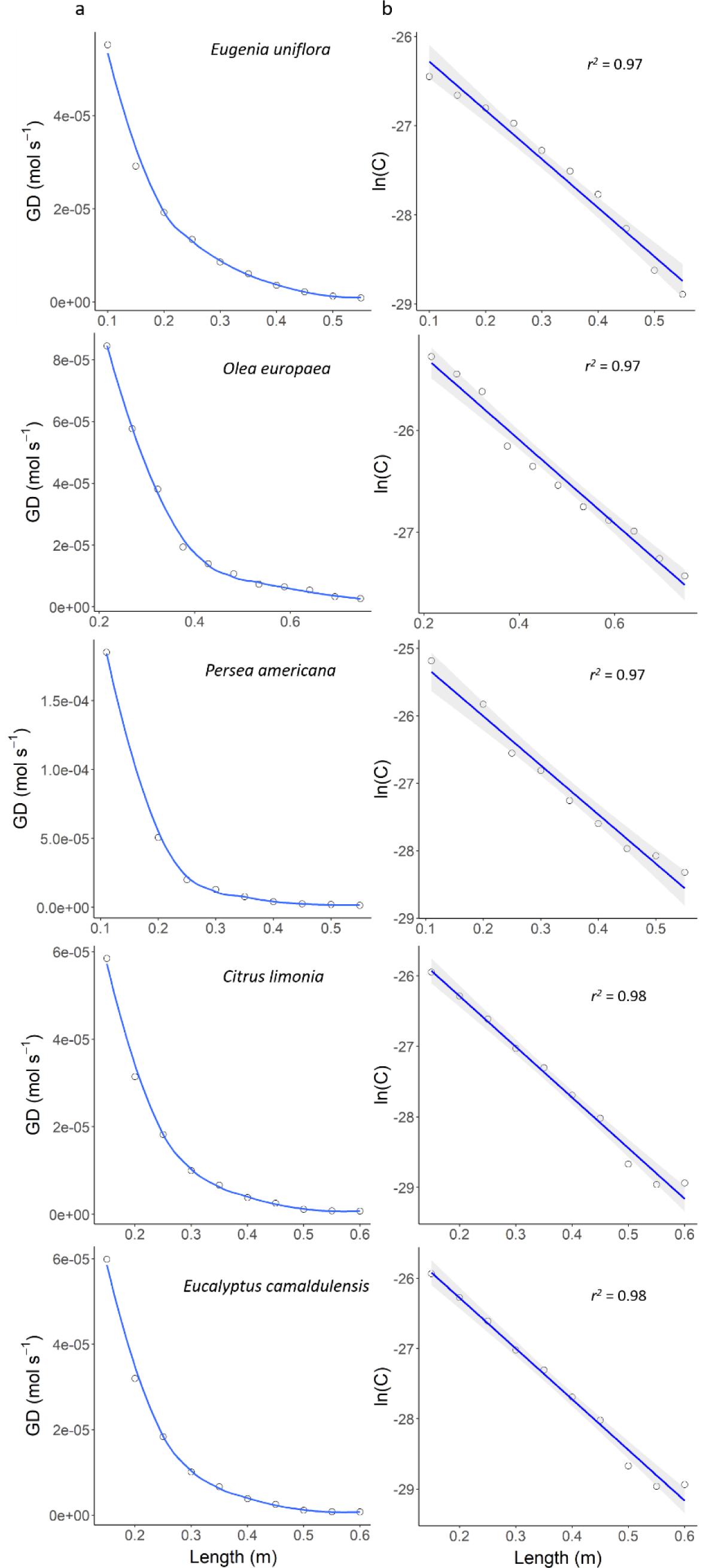
Examples of gas discharged per second (GD) and the linear relationship between the natural logarithm of gas conductivity (ln(C)) as functions of the stem segment length in *E. uniflora, O. europaea, P. americana C. limonia* and *E. camaldulensis*. All measurements are shown in Fig. S2.

The probability distribution functions (P_x_) for all species and stems are shown in Fig. 5, with the mean vessel length (L_mean_) varying from 0.19 ± 0.02 m (mean ± SD) in *Citrus limonia* to 0.45 ± 0.13 m in *Olea europaea*. Considering the species tested, the most common vessel length (L_mode_) varied from 0.10 ± 0.01 to 0.23 ± 0.06 m.

**Fig. 5.**
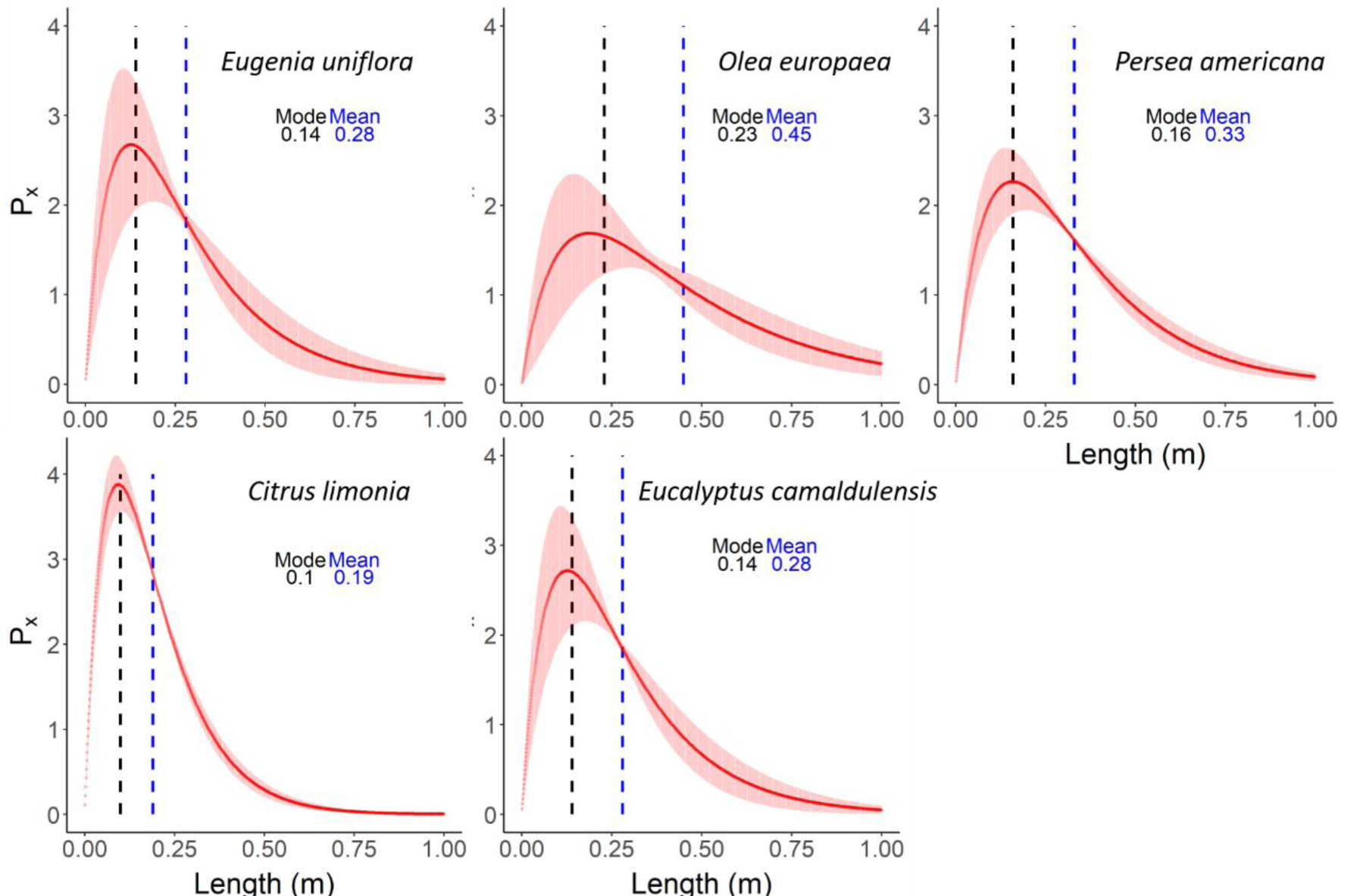
Probability distribution (Px, mean ± SD) as a function of the stem segment length in *E. uniflora, O. europaea, P. americana C. limonia* and *E. camaldulensis*. Mode (black dashed line) and mean (blue dashed line) values for vessel length estimations are shown.

A vacuum reservoir volume of 100 mL was adequate for measuring the gas flow with precision for all species, whereas 3 mL was lower than the recommended (based on Fig. 3) for measuring the gas flow in *Eucalyptus camaldulensis* and *Citrus limonia* (Fig. 6).

**Fig. 6.**
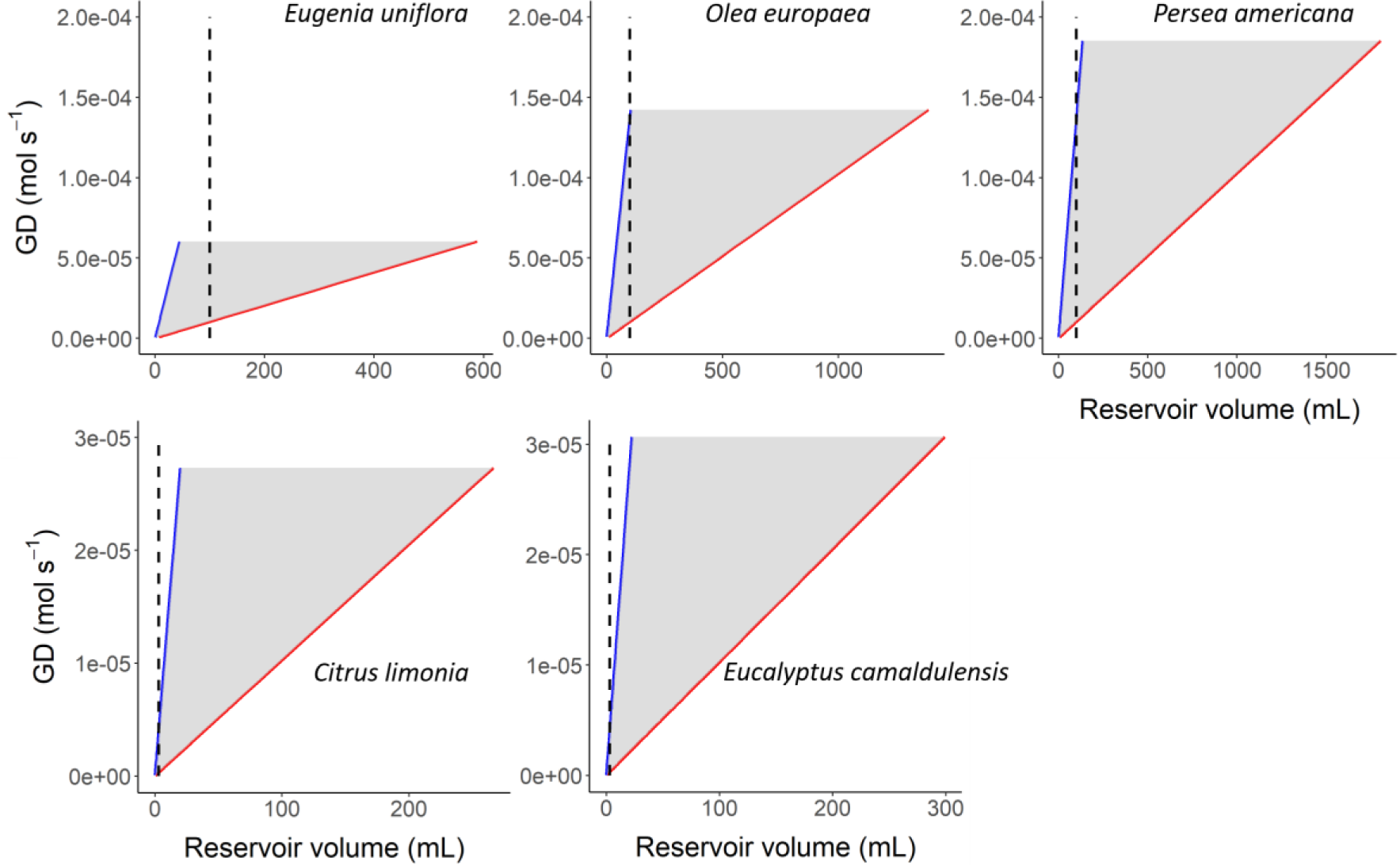
Minimum (blue) and maximum (red) reservoir volume considering all measurements of gas discharged (GD) by species. Vertical dashed lines represent the reservoir of 100 mL for *E. uniflora, O. europaea* and *P. americana*, and 3 mL for *E. camaldulensis* and *C. limonia*. Note that 100 mL is mostly within the limits of the minimum and maximum reservoir volume estimated (grey area).

During the measuring time (150 seconds), the gas conductivity showed some instability before 50 seconds when evaluating *Eugenia uniflora, Olea europaea* and *Persea americana*. In fact, the gas conductivity reached stability for all stems and species at the end of the measuring time (the last 30 seconds), varying less than 5% at the exponential increase.

## 4 Discussion

The gas conductivity values measured in our study were similar to the ones reported when using the conventional gas-conductivity method (Cohen et al. 2003; Wang et al. 2014; Pan et al. 2015), with a high correlation between the natural log of gas conductivity and stem segment length (*r2* > 0.97; Figs. 3 and 4) compared to previous reports [*r2* > 0.9 (Cohen et al. 2003); *r2* > 0.93 (Pan et al. 2015); *r2* > 0.93 (Wang et al. 2014)]. Despite applying a sub-atmospheric pressure with the Pneumatron, the exponential curves obtained were equivalent to those taken with the original gas-conductivity method, which induced flow through xylem by applying pressures above atmospheric pressure (Cohen et al. 2003).

In relation to the conventional gas-conductivity method, automated measurements are the main advantage of using the Pneumatron, which speeds up the data collection and allows multiple simultaneous measurements. We believe this is a significant improvement, especially if we consider the time needed for the silicone-injection method (Wheeler et al. 2005), which can easily surpass three weeks for ten samples. Pan et al. (2015) reported the measurement of six samples per day using the conventional gas-conductivity method, whereas we evaluated at least twice as many samples per day when using only one Pneumatron. In fact, one user can operate five to ten Pneumatrons simultaneously and, thus, the productivity could be multiplied easily. The Pneumatron is a very low-cost device based on an open-source hardware and software (Pereira et al. 2020). For this reason, the construction of many devices is accessible for most laboratories. Its enhanced capacity for collecting data is followed by simple analysis using the raw data file, with the main results being easily obtained with the R-script described here (Supplementary Material). The convenience to collect and to analyze data is important to allow pioneering studies with a large number of samples such as interbranch or interspecific comparisons, especially in highly diverse plant communities, along with measurements of other anatomical and hydraulic traits. Also, this method may be used for studies on hydraulic segmentation (Tyree and Zimmermann 2002; Levionnois et al. 2020), by estimating vessel length distribution for different plant organ connections, such as across nodes, near side branches, stem-petiole transitions, fruit pedicels, or petiole-leaf blade transitions.

Previous reports already proved the reliability of pneumatic methods for studying stem vessel length distribution (Cohen et al. 2003; Wang et al. 2014), and a comprehensive analysis of this technique is available (Cai and Tyree 2014; Pan et al. 2015). However, the results obtained with the silicone-injection and gas conductivity methods are not directly comparable. Results obtained with the gas-conductivity method are hydraulically weighted (Pan et al. 2015) and should be preferred when studying hydraulic traits, such as hydraulic resistivity of the vessel ends, or the effect of open vessels on measurements of embolism vulnerability. However, one disadvantage is that additional anatomical measurements are needed if vessel diameter and frequency are required, whereas with the silicone-injection method this is simultaneously analyzed.

The reservoir volume is an important aspect to guarantee measurement precision with the Pneumatron, which must be adequate for the gas flow and may vary among species. Although the reservoir volume that should be used may vary widely, 100 mL was suitable for most of the gas conductivity measurements (Fig. 6), and it is a reasonable starting value considering the variability found within and among plant species and samples. A smaller volume of 3 mL prevented the detection of high gas conductivity, as noticed for *Citrus limonia* and *Eucalyptus camaldulensis* (Fig. 6). Even though, the VLD estimation for these species was possible because the exponential gas conductivity increased in long stem lengths. In fact, it is recommended to use only the last three-fourth of the x-axis domain (stem length), in which gas conductivity increases exponentially due to a high number of wide vessels becoming cut open (Pan et al. 2015). As a general rule, a large reservoir volume is needed when measuring high gas flow in short stem segments. To test if the reservoir volume is appropriate, one can estimate the minimum reservoir volume by dividing the measured gas flow (GD) by 1.36759E-06, which is the slope for the relationship between the reservoir volume and GD during three seconds in which the pressure changed from 40 to 50 kPa. Similarly, the maximum reservoir volume can be obtained by dividing the measured GD by 1.02569E-07, which is the slope of the relationship between the reservoir volume and GD for 0.25 kPa of pressure change (pressure sensor linearity). Considering the equations shown in Fig. 3, users can check in a practical way how to adjust the reservoir volume for new measurements and other species.

## Supporting information

Supplementary figures and codes

## Acknowledgments

The authors acknowledge the São Paulo Research Foundation (FAPESP, Brazil) for the research grant (E.C.M, R.V.R., L.P. and M.T.M., Grant 2019/15276-8), fellowship (L.P. and R.V.R., Grant 2017/14075-3) and scholarship (M.T.M. and R.V.R., Grant 2018/09834-5). E.C.M. and R.V.R. are fellows of the National Council for Scientific and Technological Development (CNPq, Brazil). S.J., L.K., and X.G. acknowledge financial support from the German Research Foundation (DFG, project nr. 383393940 and 410768178).

