## Supplementary figures and codes for "A semi-automated method for measuring xylem vessel length distribution"

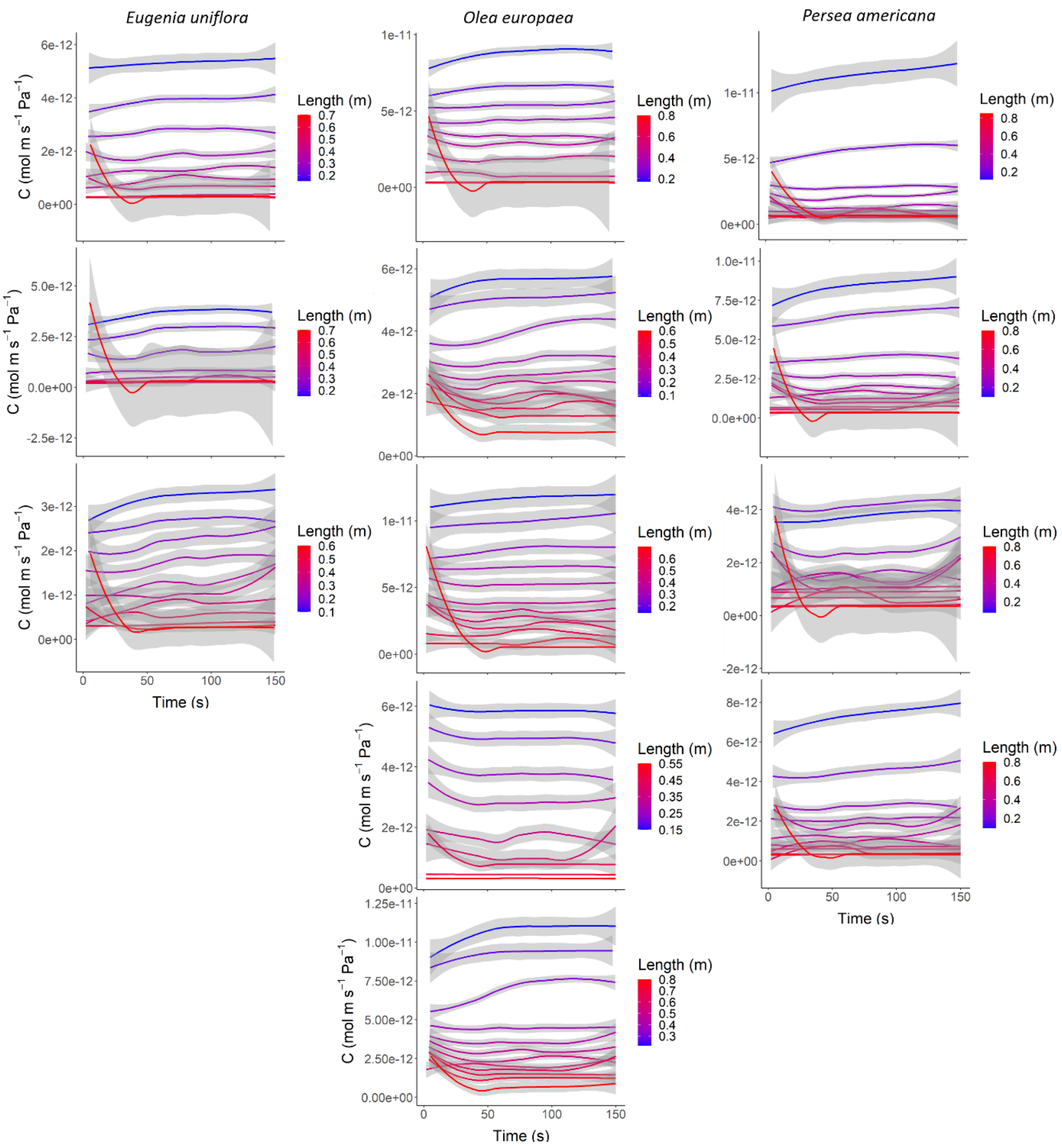


**Fig. S1** - Gas conductivity (C) during 150 seconds as affected by the length of stem segments taken from mature field-trees of *E. uniflora*, *O. europaea* and *P. americana*. Three to five samples are shown for each species.

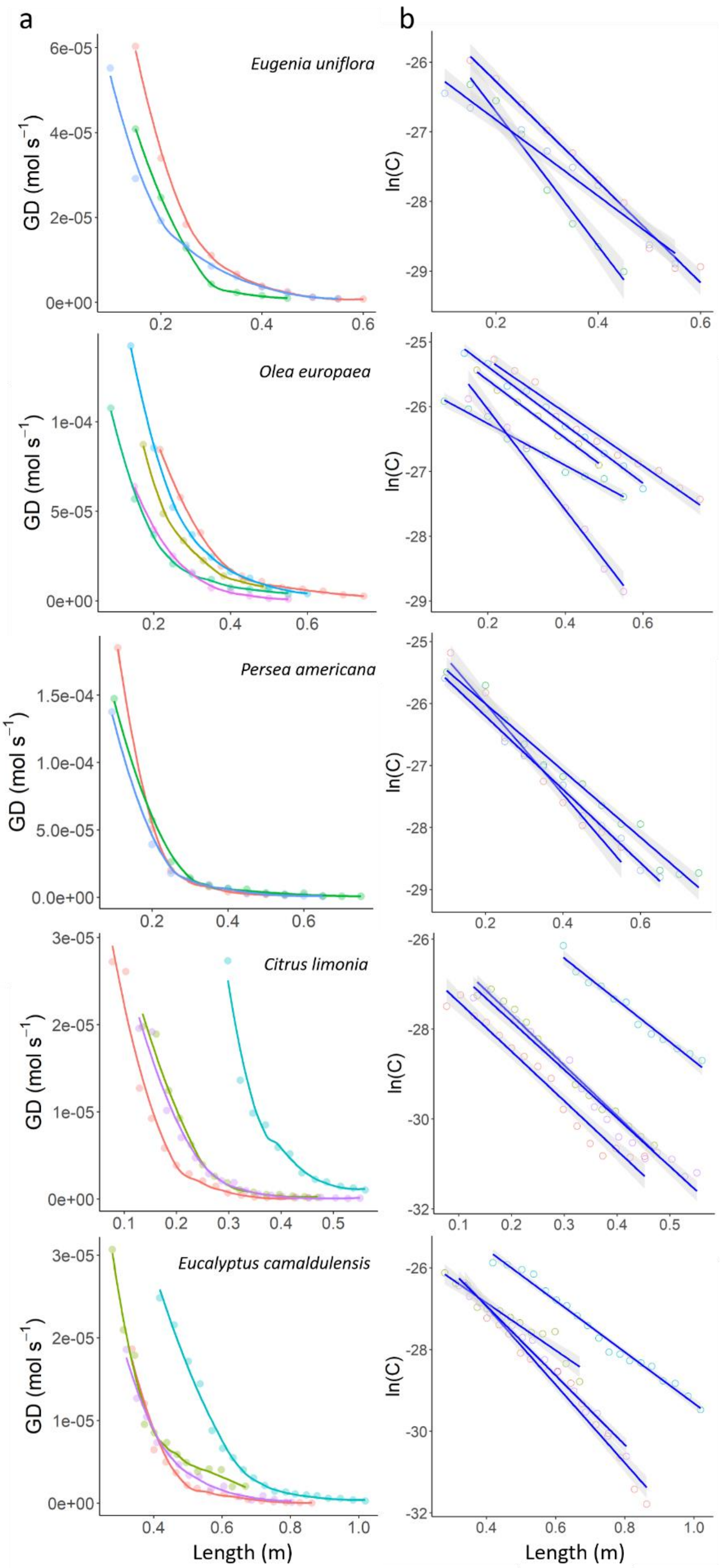


**Fig. S2** - Gas flow (a) and the natural logarithm of gas conductivity (b) as functions of the branch segment length. Each color represents different stems for a given species.  In (b), all repetitions presented $r^2 > 0.97$.

**Pneumatron software for data sampling**

```
// VESSEL LENGTH DISTRIBUTION MEASUREMENT
// A semi-automated method for measuring xylem vessel length distribution
// Luciano Pereira et al. (2020)
// Contact:

#include "RTClib.h"
#include <SPI.h>
#include <SD.h>
#include <Wire.h>
#include <Adafruit_ADS1015.h>
Adafruit_ADS1115 ads(0x48);

int Pump = 6;
int Solenoid = 5;
int Pino_CS = 10;
int buttonState = 0;
const int buttonPin = 7;

long time_1 = 1;
long time_2 = 1;
long measure = 1;

float pressure = 0.0;

RTC_DS1307 rtc;
File file;

void setup() {
  Serial.begin(9600);
  ads.begin();
  ads.setGain(GAIN_EIGHT);

  pinMode(6, OUTPUT);
  pinMode(5, OUTPUT);
  pinMode(buttonPin, INPUT);

  Serial.println("Data logger com modulo PN532");
  Serial.println();
  Serial.println("starting SD...");

  if (!SD.begin(Pino_CS))
  {
    Serial.println("error to start SD!");
    return;
  }

  Serial.println("SD card OK");
  Serial.println();

  if (! rtc.begin())
  {
    Serial.println("RTC not found!");
    while (1);
  }

  if (! rtc.isrunning())
  {
```

```
    Serial.println("RTC is not working");
    rtc.adjust(DateTime(F(__DATE__), F(__TIME__)));
  }

  rtc.adjust(DateTime(F(__DATE__), F(__TIME__)));

  file = SD.open("log.txt", FILE_WRITE);
  file.print("date");
  file.print(' ');
  file.print("hour");
  file.print(' ');
  file.print("sequence");
  file.print(' ');
  file.print("measure");
  file.print(' ');
  file.print("time");
  file.print(' ');
  file.print("pressure");
  file.println("");
  file.close();
}

void loop() {
  buttonState = digitalRead(buttonPin);
  if (buttonState == HIGH)
  {
    for(int i =0; i<300; i++)
    {
      float pressure = ads.readADC_Differential_0_1();
      pressure = ((pressure)*(0.0183)+0.297);

      if (pressure > 60) // you can change the pressure here (kPa)
      {
        digitalWrite(Pump, LOW);
        digitalWrite(Solenoid, LOW);
      }
      else if (pressure < 50) // you can change the pressure here (kPa)
      {
        digitalWrite(Pump, HIGH);
        digitalWrite(Solenoid, HIGH);
      }

      file = SD.open("log.txt", FILE_WRITE);
      DateTime now = rtc.now();
      Serial.print(now.day() < 10 ? "0" : "");
      file.print(now.day(), DEC);
      file.print('/');
      file.print(now.month() < 10 ? "0" : "");
      file.print(now.month(), DEC);
      file.print('/');
      file.print(now.year(), DEC);
      file.print(' ');
      file.print(now.hour() < 10 ? "0" : "");
      file.print(now.hour(), DEC);
      file.print(':');
      file.print(now.minute() < 10 ? "0" : "");
      file.print(now.minute(), DEC);
      file.print(':');
      file.print(now.second() < 10 ? "0" : "");
      file.print(now.second(), DEC);
```

```
            file.print(' ');
            file.print(time_1);
            file.print(' ');
            file.print(measure);
            file.print(' ');
            file.print(time_2);
            file.print(' ');
            file.print(pressure, 5);
            file.println("");
            file.close();

            delay(500);
            time_1++;
            time_2++;
          }
          digitalWrite(Pump, LOW);
          digitalWrite(Solenoid, LOW);
          measure++;
          time_2 = 1;
          delay(500);
        }
      }
```

**R-script for data analysis**

```
####### VESSEL LENGTH DISTRIBUTION ANALYSIS ###################################
####### A semi-automated method for measuring xylem vessel length distribution #################
####### Luciano Pereira et al. (2020) #####################################################
####### Contact: #################################################
library(dplyr)
library(ggplot2)
library(ggpubr)

# Two files needed:
# 1 - The raw data file from Pneumatron
# 2 - two columns .csv file indicating the (1) measure, (2) length

setwd("C:/Users/…") # select your folder
raw <- read.delim("log.txt", header = T, sep = ' ', as.is = T) # select the raw data file
id <- read.csv("species_id.csv", header = T, sep = ',', as.is = T) # select the file which indicates the measure and length

## define experimental conditions
reservoir <- 100 # in mL
atm <- 101325 # atmospheric pressure in Pa
set_r2 <- 0.97 # insert the minimum r2 considered for the slope
Vr <- reservoir * 10^-6 # reservoir in m^3
R <- 8.31446213 # gas constant
t <- 293.15 # in K
###################################

## correct data format and insert new variables
raw$sequence <- seq(1, nrow(raw))
raw1 <- merge(raw, id, by = "measure")
raw1$time <- as.numeric(raw1$time)
raw1$pressure <- as.numeric(raw1$pressure)
raw1$length <- as.numeric(raw1$length) / 100 # length per measure in m
raw1$p_abs <- atm - (raw1$pressure * 1000) # absolute pressure in Pa

raw1 <- raw1[order(raw1$sequence), ]
raw1$t <- raw1$time * 0.5
raw1$p_abs_ini <- lag(raw1$p_abs, n = 3) # n=3 means interval of 1 second
raw1 <- raw1[raw1$time > 3, ] # delete lag lines NA or from another length
raw1$gd_mol <- ifelse(raw1$p_abs > raw1$p_abs_ini,
                      ((raw1$p_abs * Vr) / (R * t)) - ((raw1$p_abs_ini * Vr) / (R * t)), # Equation 1
                      NA)
raw1 <- na.omit(raw1)
##################################################

## calculate conductivity (C) according to Pan et al. (2015)
raw1$c <- (raw1$gd_mol * raw1$length) / ((atm - raw1$p_abs_ini) / 2) # Equation 2
raw1$ln <- log(raw1$c)
################################################################

## conductivity at the last 30 seconds
raw2 <- raw1[raw1$time > 240, ]
########################################

## Remove values that do not fit to r square
mean1 <- raw2 %>%
  group_by(measure) %>%
  summarise_all(funs(mean)) # calculate means
model <- lm(ln ~ length, data = raw2) # generate regression line
```

```r
r2 <- summary(model)$r.squared # calculate r squared from model
x <- 1

## remove non-exponential data points
repeat {
   mean2 <- mean1[mean1$measure > x , ]
   model <- lm(ln ~ length, data = mean2)
   r2 <- summary(model)$r.squared
   print(x)
   print(r2)
   x <- x + 1

   if (r2 > set_r2){
      break
   }
}
##############################################

## Calculate probability function
int <- summary(model)$coefficients[1]
slope <- summary(model)$coefficients[2] # λv
result <- data.frame(int, slope)
df <- data.frame("length" = seq(0, 1, by = 0.001))
df$n <- int * exp(slope * df$length) # Equation 3
df$px <- df$length * (slope^2) * exp(slope * df$length) # Equation 4
Lmode <- -1 / slope # Equation 5
Lmean <- -2 / slope # Equation 6
##################################


#### graphs
## Graph from Fig. S2
fig_slope <- ggplot(data = mean2, aes(x = length, y = ln)) +
  geom_point(shape = 21, size = 3) +
  geom_smooth(method = lm , color = "blue", fill = "lightgray", se = TRUE, size = 1) +
  scale_x_continuous("Length (m)") +
  scale_y_continuous("ln(C)") +
  theme_bw() +
  theme(panel.border = element_blank(),
        panel.grid.major = element_blank(),
        panel.grid.minor = element_blank(),
        axis.line = element_line(colour = "black"),
        aspect.ratio = 1) +
  stat_cor(label.x = 0.3, label.y = -26.5) +
  stat_regline_equation(label.x = 0.3, label.y = -26.3)

fig_slope

## Graph from Fig. 5
fig_px <- ggplot(data = df, aes(x = length, y = px)) +
  geom_line(size = 1, colour = "red") +
  scale_x_continuous("Length (m)") +
  scale_y_continuous("Px") +
  theme_bw() +
  theme(panel.border = element_blank(),
        panel.grid.major = element_blank(),
        panel.grid.minor = element_blank(),
        axis.line = element_line(colour = "black"),
        aspect.ratio = 1) + # formato final
  theme(axis.title.x = element_text(vjust = -1.0),
```

```
      axis.title.y = element_text(vjust = +1.0))+
  annotate("segment", y = 0, yend = 3, x =   Lmode, xend = Lmode,
           colour = "black", size = 1, linetype = "dashed") +
  annotate(geom = "text", x = 0.7, y = 2.5, label = "Mode", color = "black")+
  annotate(geom = "text", x = 0.7, y = 2.4, label = format(round(Lmode, 2), nsmall = 2), color = "black") +
  annotate("segment", y = 0, yend = 3, x =   Lmean, xend = Lmean,
           colour = "blue", size = 1, linetype = "dashed") +
  annotate(geom = "text", x = 0.8, y = 2.5, label = "Mean", color = "blue") +
  annotate(geom = "text", x = 0.8, y = 2.4, label = format(round(Lmean, 2), nsmall = 2), color = "blue")

fig_px

## graph from Fig. S1
fig_cond <- ggplot(data = raw1, aes(x = t, y = c, group = length, color = length)) +
  geom_smooth(method = "loess", se = TRUE) +
  scale_x_continuous("Time (s)") +
  scale_y_continuous(bquote('C (mol' ~ m ~ s^-1 ~ Pa^-1 * ')')) +
  theme_bw() +
  theme(panel.border = element_blank(),
      panel.grid.major = element_blank(),
      panel.grid.minor = element_blank(),
      axis.line = element_line(colour = "black"),
      aspect.ratio = 1) +
      scale_colour_gradient(name = "Length (m)", low = "blue", high = "red")

fig_cond

ggsave("fig_slope.png", fig_slope, width = 4.5, height = 4.5, units = "in")
ggsave("fig_px.png", fig_px, width = 4.5, height = 4.5, units = "in")
ggsave("fig_n.png", fig_n, width = 4.5, height = 4.5, units = "in")
ggsave("fig_cond.png", fig_cond, width = 4.5, height = 4.5, units = "in")
```
